# Robust brain complexity measurement using T2* informed multivariate sample entropy from multi-echo BOLD fMRI

**DOI:** 10.64898/2025.11.29.691196

**Authors:** Ning Wang, Mingyan Wu, Weihuan Fang, Jiawen Sun, Liwen Zhang, Naying He, Fuhua Yan, Kui Ying, Xingfeng Shao

## Abstract

Sample entropy is a useful tool to analyze resting-state fMRI data and evaluate the complexity of neural activity. However, sample entropy computed from conventional BOLD signals is not highly resistant to artifacts. In contrast, T2* may be more sensitive to certain neural-related changes, and directly incorporating T2* information enables the detection of features that are often lost in BOLD-only sample entropy due to noise influences. To further improve the robustness and reliability of complexity estimation, we introduce a multivariate sample entropy approach that incorporates T2* signals derived from multi-echo BOLD signal. We applied this method to compare a cohort of 23 elder adults and 12 younger adults, and observed significant group differences in several brain regions. Notably, the proposed method yielded more pronounced group differences than conventional sample entropy computed without T2* information. Moreover, in regions with reduced neural activity due to ischemia, as well as in ventricular areas, our approach demonstrated superior performance. These results indicate that the proposed method enhances the ability to characterize the complexity of neural activity.

## Introduction

Recent studies have shown that the temporal complexity of blood-oxygen-level-dependent functional magnetic resonance imaging (BOLD-fMRI) signals can reflect the brain’s capacity for information processing.^[1][2][3]^

Task-based fMRI measures the BOLD signal differences between the resting state and when brain stimulated by a specific task to identify associated brain regions ^[4]^. In contrast, resting-state fMRI focuses on spontaneous low-frequency fluctuations in the brain ^[5]^. It is commonly used to calculate functional connectivity, which is defined as the temporal correlation between two brain regions. Through this approach, resting-state networks can be established, revealing default patterns of interaction in the brain and providing a better understanding of intrinsic brain activity ^[6]^. However, functional connectivity only reflects the inter spital connection between different brain regions and cannot capture the voxel-wise temporal dynamics of the time series ^[7]^.

Recently various information-theoretic methods have been developed to characterize the temporal complexity and dynamics of physiological systems using rs-fMRI ^[8,9]^. ‘Entropy’ is a measure of a system’s chaos level, which is also a powerful tool to analyze brain complexity. Entropy of BOLD signals quantify the likelihood of new pattern generation to evaluate the complexity of the human brain, and is highly associated with consciousness, brain aging, and cognitive function in neurodegenerative diseases such as AD. ^[10][11].[12]^ Studies have also shown that physiological aging in certain regions of the human brain can lead to cortical atrophy and reduced neurotransmitter levels.^[13]^ In addition, lifespan trajectories of brain functional complexity have been constructed, demonstrating that entropy is a sensitive metric capable of capturing age-related differences in brain function.^[14]^

Several approaches have been developed to quantify brain entropy, including differential entropy and permutation entropy. Approximate entropy, proposed by Steve Pincus in 1995, characterizes the regularity and complexity of time-series data. Sample entropy was later introduced as an improvement over approximate entropy, reducing its bias and providing more reliable estimates, particularly for short time series ^[15-18]^. However, the applications of existing methods are still limited due to their insensitivity to neural signals in the presence of noise^[19]^.

The BOLD signal is influenced by multiple physiological factors, including cerebral blood flow (CBF), cerebral blood volume (CBV), and the cerebral metabolic rate of oxygen (CMRO_2_). One of the major contributors to the BOLD signal is the difference in magnetic properties between hemoglobin and deoxyhemoglobin, and is highly associate with the high-frequency neuronal activity this is not right, BOLD signal comes from neurovascular coupling effect, fresh blood carrying extra hemoglobin leading to increased BOLD signal^[20]^. Thus, changes in T^2^* induced by hemoglobin-level fluctuations may provide a more direct indicator of underlying neural activities^[21][22]^.

To improve the reliability of the BOLD signal, multi-echo BOLD sequences have been used to enhance contrast and reduce physiological artifacts ^[23]^. Compared to the conventional BOLD acquisition, multi-echo BOLD has several advantages. TE-dependent non-BOLD components can be separated from BOLD-like components^[24]^. However, brain entropy always only evaluates patterns in the BOLD signals, whereas BOLD T2* signal can be more sensitive to subtle changes in oxygenation and brain activities^[25]^. Directly using the T2* signal and placing more emphasis on its dynamics may contribute to the analysis of neural activity. In addition, SNR in ultra-high field (UHF) MRI increases superlinearly with field strength^[26,27]^, which can further improve the reliability of the brain entropy measurements ^[28]^.

The goal of this study is to propose an innovative quantification method to obtain robust brain complexity measurement. We calculated brain entropy from multi-echo BOLD signals using a multivariate sample entropy approach incorporating T2* signals. Theoretical improvement of the proposed method compared to conventional sample entropy method was demonstrated by theoretical simulations and experiments at two different field strengths. Age-related changes in neural complexity was studied in young and elderly populations, and alterations neural activity in ischemic lesion was further demonstrated in a patient with middle cerebral artery (MCA) stenosis.

## Methods

### MRI experiments

Experiments were conducted on a 5T MRI scanner (United Imaging, uMR Jupiter) using a 2Tx/48Rx head coil, multi-echo BOLD signals were acquired with the following parameters: spatial resolution = 2.2 mm^3^ isotropic; TE = [9.0, 23.8, 38.6, 53.4] ms; TR = 1.8 s; 69 slices; 210 repetitions; total scan time = 6 minutes 6 seconds. Data preprocessing steps included temporal realignment, slice timing correction, linear detrending and using a low-pass filter of 0.2Hz, and these steps were implemented using SPM and CONN^[29]^. The first ten volumes before reaching steady state were discarded. Images were spatially normalized to the Montreal Neurological Institute (MNI) template space for group-level analysis.

To evaluate whether higher magnetic field strength leads to improved results, multi-echo BOLD fMRI signals were also acquired at 3T with following parameters: TR = 1.35 s, TEs = [10.8, 32.2, 53.6, 74.0] ms, and spatial resolution = 2.5 mm^3^, 48 slices, 210 measurements, total time = 4minutes 40 seconds,.

A multi-delay arterial spin labeling (ASL) with six post-labeling delays (PLDs) was also acquired from the patient with MCA stenosis, cerebral blood flow (CBF) and arterial transit time (ATT) maps were calculated according to Shao et al^[30][31]^. Ischemic lesion was identified as regions with significant reduced CBF and prolonged ATT.

### Signal processing

The combined BOLD signal was calculated according to the following equations:

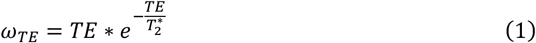

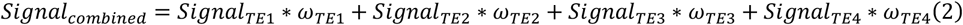

According to Eqs. [1,2], the T2*-weighted decay is employed to combine the echoes via weighted averaging. The resulting combined data generally exhibit higher signal-to-noise ratio (SNR) and greater statistical power ^[24]^.

T2* can be estimated from multi-echo BOLD signals using nonlinear least-squares fitting of Eq. [3]:

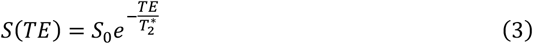

As illustrated in Fig. 1, we propose a novel method to calculate multivariate sample entropy (MVSE) by combining both z-score normalized BOLD magnitude and T2* signals.

**Figure 1.**
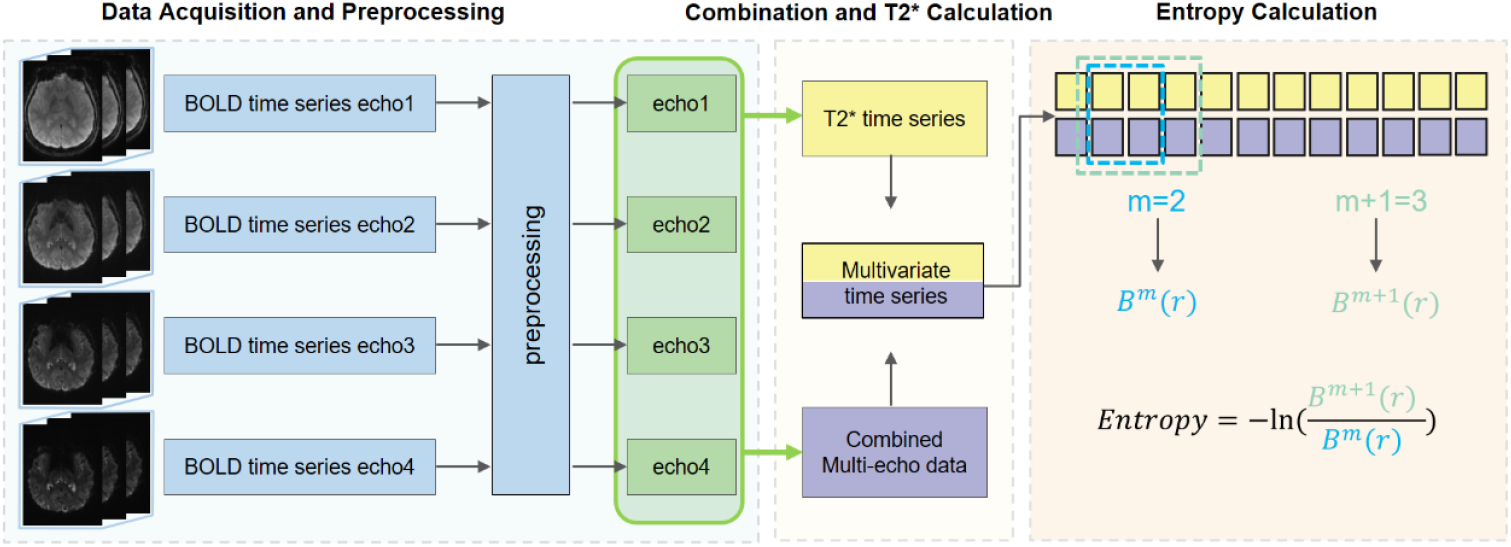
The workflow of computing multivariate sample entropy: T2* values were first calculated from multi-echo BOLD signals, then jointly computed with combined BOLD signals.

To ensure balanced contributions to the MVSE the standard deviation was calculated jointly from the T2*-weighted and BOLD signals. Subsequences (a short continuous segment) of the dynamic signals with a Chebyshev distance less than 0.6 times this standard deviation were considered similar. Two signal sequences, u(i) and u(j), were considered matched only if the distance criterion was satisfied in both modalities.

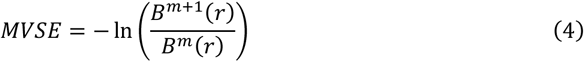

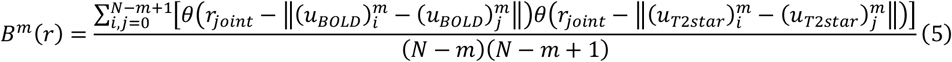

B^m^(r) refers to the frequency of occurrence that two subsequences will match for m points with a tolerance of r. 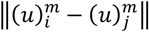 represents the distance between two sequences, θ stands for the Heaviside step function. This dual-criterion matching indicates stronger and more consistent neural activity. A comparison was also made with sample entropy calculated using BOLD magnitude signals or T2* only.

Multiscale analysis was performed at three different scales (scale 1, 2 and 3) to find the best scale to evaluate the complexity. During the coarse-graining process at scale τ, the sequence with length N was locally averaged to generate a new sequence of length 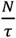, following the rule below:

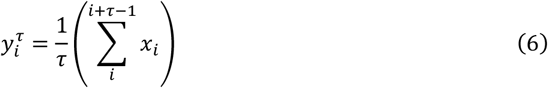

After this process, the high-frequency components of the sequence are removed, allowing the time series to be evaluated at different temporal scales.

To compare data quality between the 3 T and 5 T acquisitions, the signal-to-noise ratio (SNR) was calculated, which quantifies the quality of BOLD signal intensity across consecutive time points. SNR for each 4D fMRI dataset was computed using the following formulation ^[32,33]^:

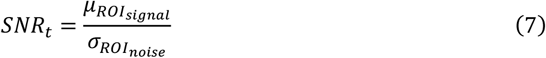

Where *μ*_*ROI_signal*_ is the average of signal in the brain mask, *σ*_*ROI_noise*_stands for the standard devison of noise area at time t.

Simulations were performed to analyze the effect of multiscale analysis. Signals with different frequencies were generated, and Gaussian noise was then added to simulate high-frequency fluctuations.

### Participants

A total of 35 healthy volunteers were recruited including 23 elderly subjects (9 males, age =67.8±5.8 years) and 12 young subjects (9 males, age = 24.4±3.0 years). One patient (male, 68 years old) with right side MCA stenosis was also included to demonstrate the brain complexity changes in ischemic regions. All participants were provided with informed consent (IRB number: RUIJIN2025-109).

### Statistical analysis

For the young and elderly groups, we used a t-test to compute the p-values for statistical significance, and Cohen’s d to compare the effect sizes across different entropy calculation methods. Functional connectivity was calculated using Pearson correlation, and a t-test was performed to analyze brain activity and assess the entropy.

## Results

The results of the multiscale analysis on the simulated data are shown in SI Figure S2 and S3. For a regular sine function in SI Figure S2, the entropy does not show a substantial decreasing trend as the scale increases. However, when noise is added to the signal, entropy decreases significantly with increasing scale, indicating that the coarse-graining process efficiently reduces the influence of noise. Similarly, a low-pass filter can partially remove high-frequency components. It has a noticeable effect when the scale is small, but as the scale increases moderately, the difference before and after filtering becomes minimal. The signals in SI Figure S3 contain two frequency components. When analyzed in the same way, applying a low-pass filter does not lead to substantial improvements at larger scales.

The results calculated from 3 T and 5 T scans from the same subject are shown in the figure 2. For both MVSE and SE, the ratio of the mean signal in gray matter to that in white matter was 0.93 at 3 T, increasing to 0.98 at 5 T. These values are consistent with the results reported by Robert X. Smith et al., 2013 ^[34]^. The contrast between gray matter and white matter indicates that more neural signal features are captured at 5 T than at 3 T. Although this does not necessarily imply that the results at 5 T are superior, this pattern is more consistent with physiological expectations, suggesting that 5 T data can better reflect neural complexity and get more reliable calculations. In addition, artifacts observed in the frontal lobe area at 3 T were not observed at 5 T.

**Figure 2.**
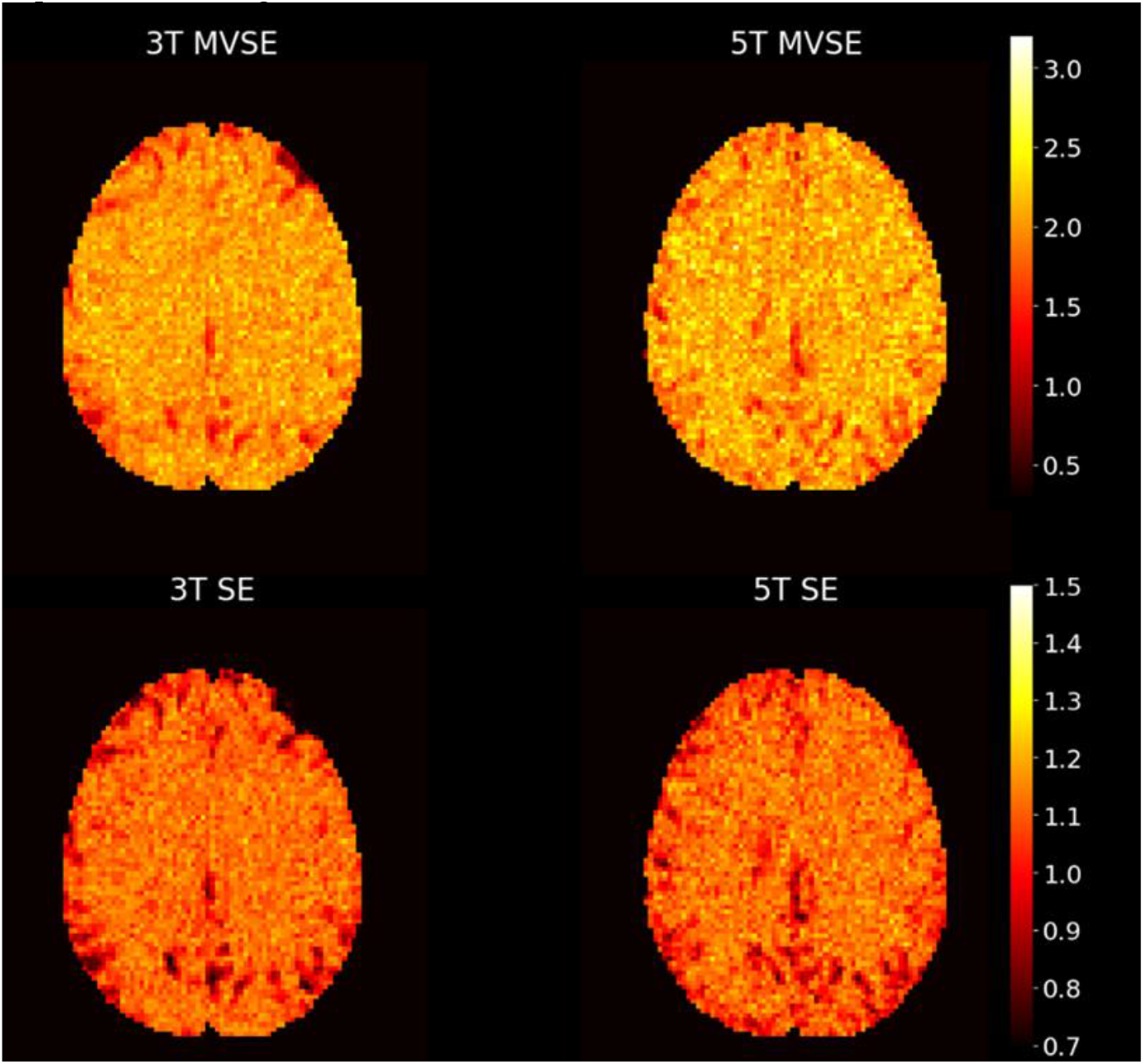
Results of four-echo BOLD signals acquired from the same subject at 3 T and 5 T. The signals were analyzed using MVSE (up) and SE down).

Figure 3 shows the group-averaged maps of the proposed MVSE for the elderly and young groups in normalized MNI space. Lower entropy can be observed in in the elderly group, particularly in the gray matter. The mean gray matter value decreased from 1.81 in the young group to 1.65 in the elderly group.

**Figure 3.**
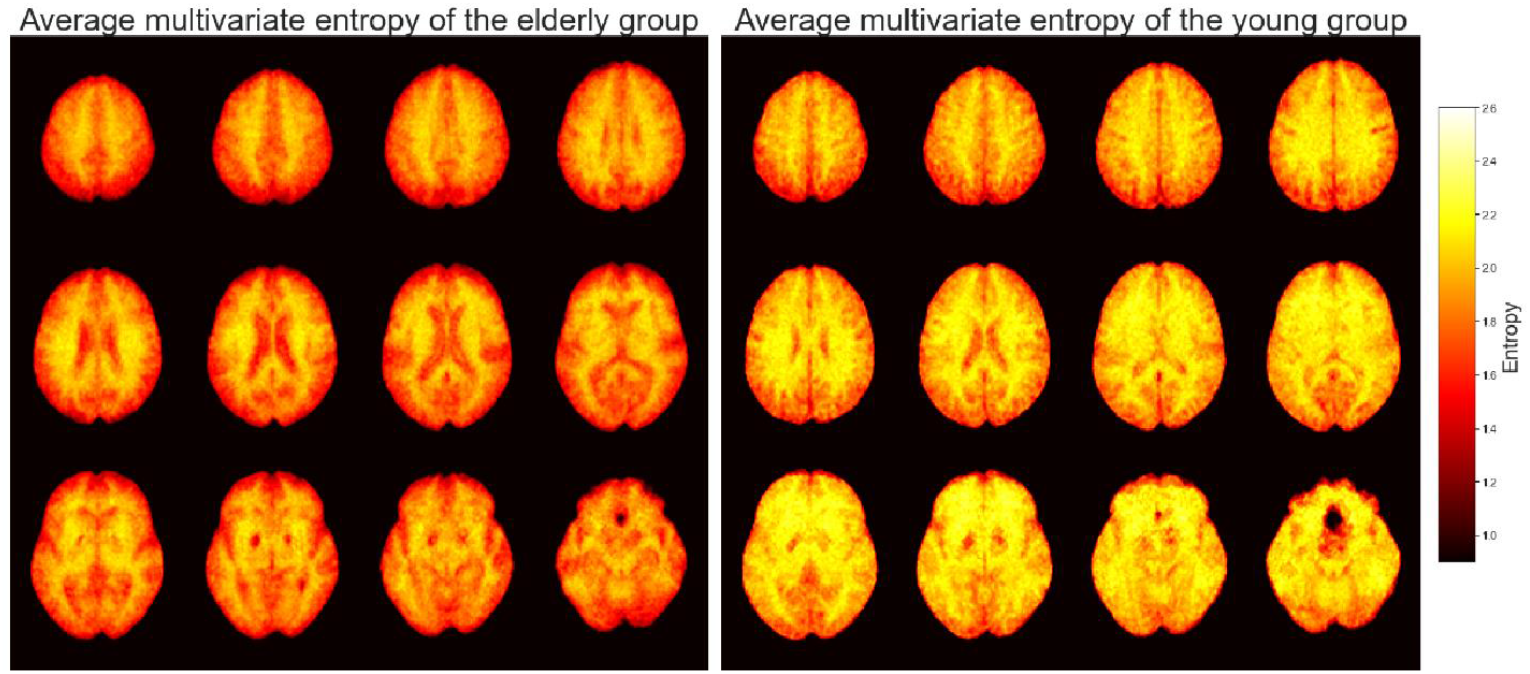
Group average of the proposed multivariate sample entropy maps from the elderly (left) and young (right) groups. Lower entropy can be observed in the aged group especially in the gray matter.

Figure 4 shows the distribution of mean entropy in different brain regions for both elderly and young groups. A significant difference was observed between the elder and younger groups, the proposed method leads to more significant inter-group differences and a larger effect size (Cohen’s d). The results demonstrate that adding T2* as an additional variable allows entropy to capture more information than using BOLD magnitude signals alone. T2* time series provide more information related to underlying neural activities for calculating entropy and assessing brain complexity.

**Figure 4.**
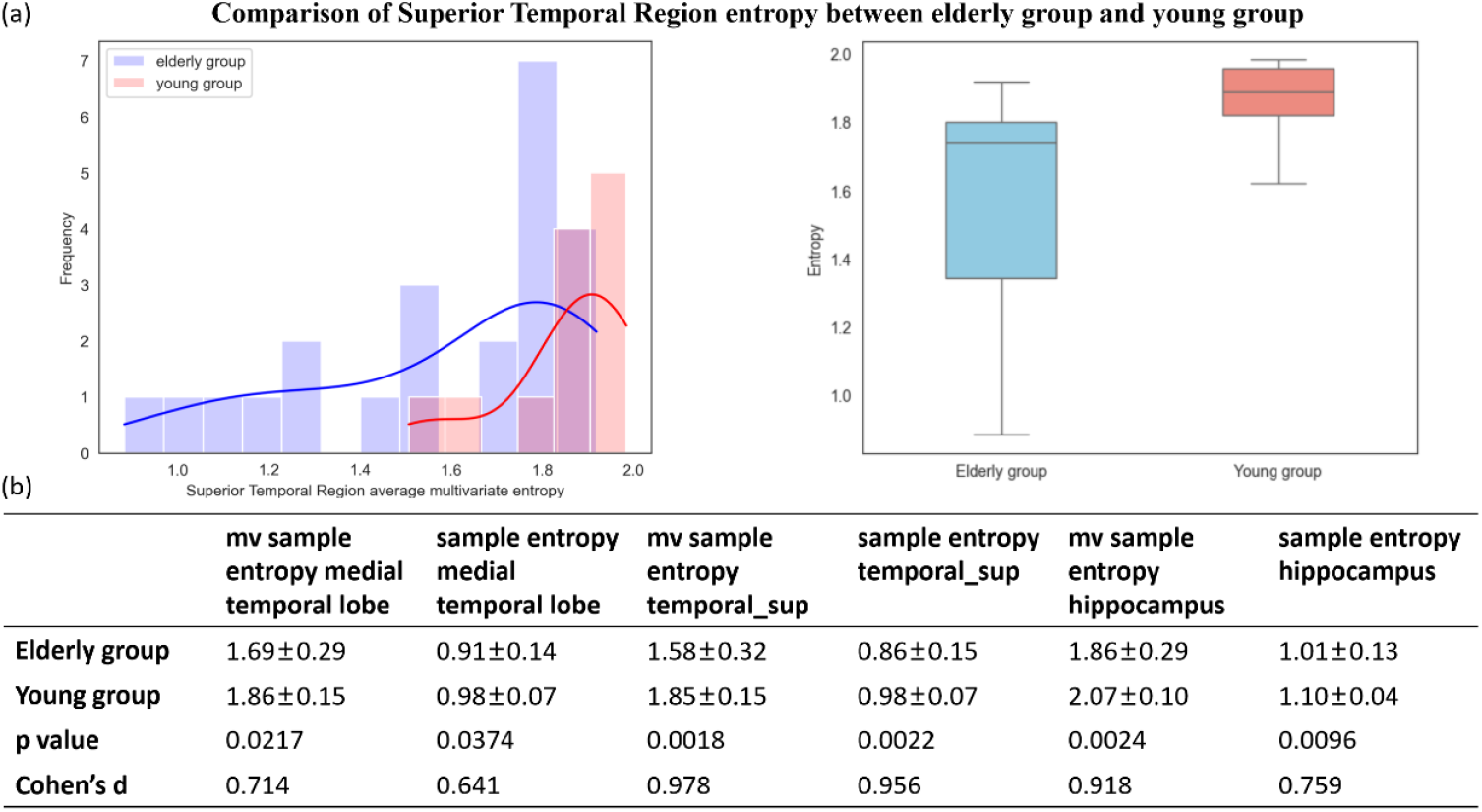
(a) Histogram and box plot of superior temporal region entropy; (b) Table of average multivariate (mv) sample entropy, conventional BOLD sample entropy between the elderly and young groups. Significant differences between the two groups can be observed in the hippocampal and superior temporal regions.

Figure 5 shows that the differences between the young and old groups become more pronounced as the scale increases. However, due to the relatively short length of the time series used in this study (200 measurements), entropy maps became nosier when larger temporal scales are applied.

**Figure 5.**
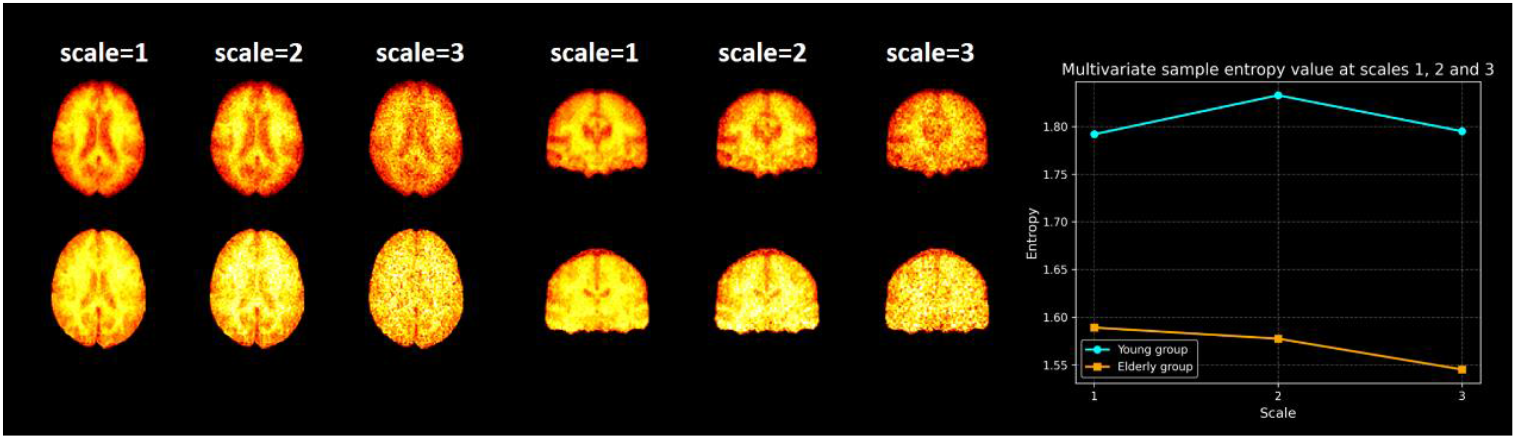
Multiscale analysis of multivariate sample entropy. From left to right: scale = 1, scale = 2, and scale = 3. The first row shows the mean entropy mapping for the elderly group, and the second row for the young group. The plots show the mean entropy values in gray matter regions for both groups across different scales.

Figure 6 (a-e) shows CBF and ATT maps calculated from multi-delay ASL, the proposed MVSE, sample entropy maps derived from BOLD magnitude signals or T2* signals from a representative individual (male, 68 years old). Ischemic region was highlighted by dashed white boxes, with potentially reduced neural activity. BOLD SE is less sensitive to such changes; however, both the T2* SE and the proposed MVSE successfully detect reduced complexity. Interestingly, we observed similar BOLD SE signals in the ventricle as compared to GM/WM regions, while neural activity should not be expected in ventricles when only CSF is presented. In contrast, both MVSE and T2* SE maps show reduced complexity suggesting improved reliability as compared to conventional BOLD SE.

**Figure 6.**
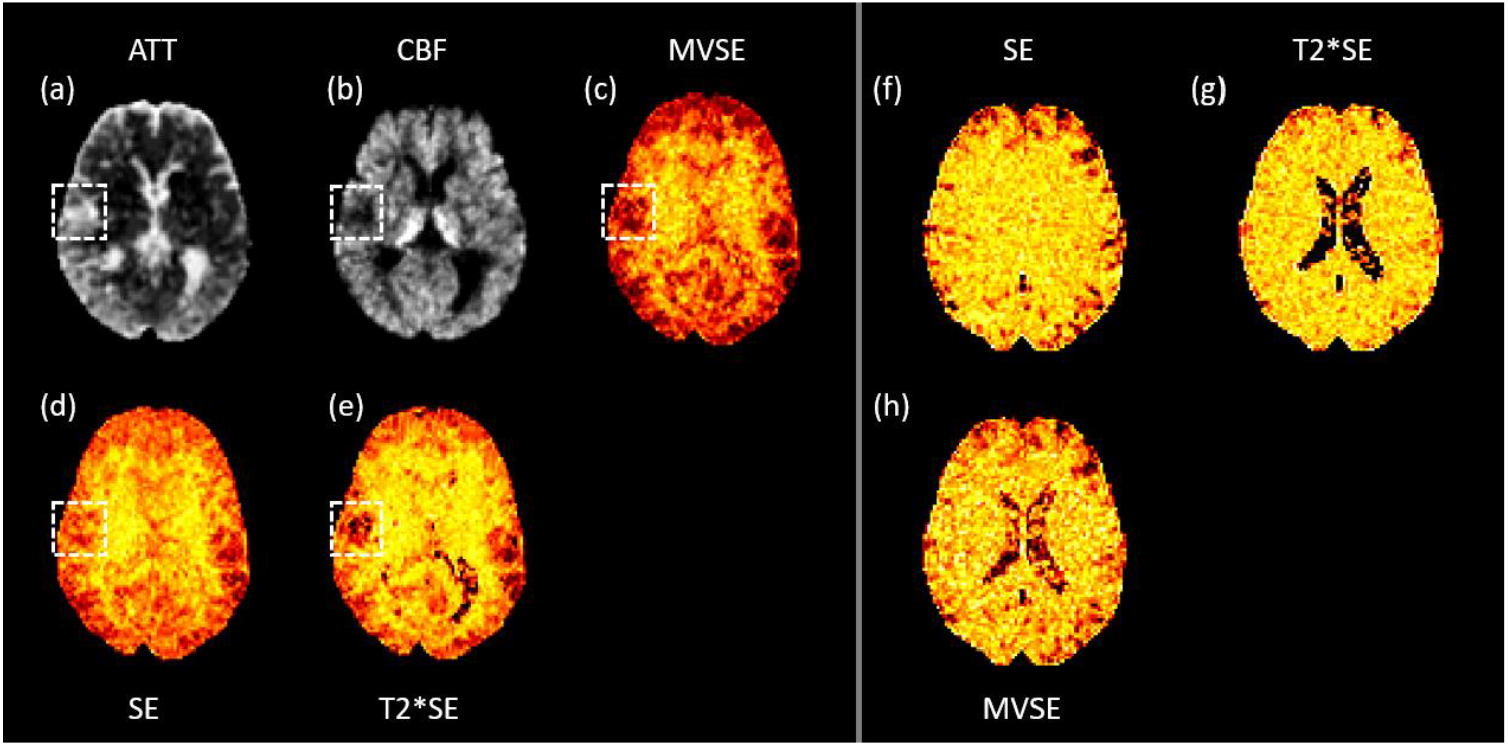
(a) Arterial transit time (ATT) map, (b) cerebral blood flow (CBF) map, (c) multivariate sample entropy, (d) sample entropy, and (e) entropy derived from T2* signal for a representative patient. (f) sample entropy (g) entropy derived from T2* signal (h) multivariate sample entropy for another representative patient.

## Discussion

In this study, we proposed a MVSE method to calculate the brain complexity. Improved robustness in complexity measurement as compared to conventional sample entropy method has been demonstrated by simulations and experiments at two different field strengths. The proposed method is also capable of detecting age-relevant variations in neural complexity, as well as reduced neural activity in regions with insufficient blood supply.

The comparison of data quality between 5 T and 3 T is presented in SI Figure S1, we observed that at 5 T, the SNR of echo 1 was not the highest. This may be due to the relatively short TE of 9 ms, which could prevent the 180° inversion pulse from fully reaching a steady state. Additionally, this effect may be related to B1 inhomogeneity.

Using the proposed method, we observed significantly lower neural complexity in elderly group (Figure 3). In particular, a clear reduction in entropy can be observed in the temporal superior and temporal lobe regions among the elderly group. This indicates that with aging, the ability of certain brain regions to generate new neural signal patterns decreases, reflecting a decline in neural activity complexity. These findings provide evidence for age-related degeneration of neural function and suggest that the proposed metric may serve as a potential biomarker to aid in diagnosis.

Existing studies have shown that CBF in the gray matter of patients with mild cognitive impairment (MCI) is significantly reduced compared with healthy controls, and this reduction is even more pronounced in individuals with Alzheimer’s disease (AD). In addition, cerebral oxygen extraction fraction (OEF) measured with BOLD MRI provides further insight into altered metabolic demands in these populations. Research using time-shift analysis has also demonstrated its utility in assessing hemodynamic lag, with ischemic stroke regions typically exhibiting delayed BOLD signal timing within the corresponding vascular territories.^[35,36]^

Ischemic strokes account for the majority of stroke cases and are a major cause of long-term disability and cognitive impairment^[37]^. Neuronal injury in these regions may arise from multiple mechanisms, including ischemia-induced neuronal loss, infarction, and vascular obstruction.

As T2* is highly sensitive to oxygenation fluctuations, reductions in neural activity in ischemic regions are more effectively captured. In contrast, in the ventricle regions, although the complexity of the BOLD signal does not show a clear decrease in entropy, the decay of BOLD signals across different echo times (i.e., the T2* signal fluctuations) can effectively capture the process of reduced neural activity. The proposed method therefore performs better in both ventricle and ischemic regions, indicating that the T2* signal is more sensitive to neural activity changes.

We also observed great within-group variability in the elderly group. This is largely attributable to the heterogeneous composition of the group, which included participants with Alzheimer’s disease (AD), mild cognitive impairment (MCI), and cognitively normal elder adults. Such heterogeneity introduces additional factors beyond age that may contribute to the observed entropy differences. Although we applied motion correction and denoising procedures, the elderly participants generally showed lower compliance during scanning compared with the younger group, which may have introduced additional uncertainty into the computed results. Additionally, in the young group, we observed a few individuals with unusually low values. This may be related to the fact that they did not fully keep their eyes open during the scan and may have entered a very light sleep state^[38]^.

Different time scales are often used to assess neural activity cycles at different frequencies. However, in this experiment, the time series is relatively short, so multiscale sample entropy does not perform well. Although sample entropy is generally suitable for short time series, at scale = 4, the effective sequence length is only 50, which introduces substantial noise in the calculation. Currently, the multiscale analysis applies a unified coarse-graining process for the dual-channel data. However, T2* signals and multi-echo BOLD signals may have different fluctuation frequencies during the resting state. Therefore, a future improvement could be to implement a dual-input multiscale analysis, performing coarse-graining separately at different scales for each channel.

In SI Figure S1, it can be observed that SNR increases with longer echo times, indicating reduced image quality at later echoes. In contrast, the 5T acquisition shows consistently higher SNR values, reflecting improved image quality to 3T. This demonstrates that ultra-high-field imaging provides clearer and more stable BOLD signals. Furthermore, combining multiple echoes further enhances data quality by leveraging complementary information across TEs.

SI Figure S2 and SI Figure S3 shows the coarse-graining process is conceptually similar to applying a low-pass filter: both reduce the high-frequency components of the time series, smoothing the signal. In this process, noise in the signal may be removed, allowing the underlying neural activity to be more clearly reflected.

The functional connectivity was also calculated to further evaluate the performance and reliability of the entropy measure. In SI Figure S4, connectivity analysis was performed across 45 regions of interest defined by the AAL atlas. The average connectivity strength between regions was computed using Pearson correlation coefficients. Connectivity loss was observed in several brain regions, and certain region pairs exhibited substantial reductions in connectivity, as summarized in the table. Notably, these regions also demonstrated large differences in multivariate entropy, supporting the sensitivity of the proposed method.

There are several limitations of this study. The observed decrease in entropy in the elderly group may not be driven by a single factor. It could be influenced not only by aging itself, but also by underlying conditions such as Alzheimer’s disease (AD), mild cognitive impairment (MCI), or other age-related pathologies. Therefore, it is difficult to determine which factor contributes more to the difference observed between young group and the elderly group.

More data from strictly screened healthy elderly participants are needed to further clarify which factor plays the most important role in the entropy reduction, to understand the underlying mechanisms. In this way, entropy may also serve as a tool to help identify the underlying factors contributing to neural activity decline, including whether amyloid-β or tau pathology plays a role in driving this decrease.

The calculation of entropy is influenced by scanning parameters, such as different TR values, which may require adjusting the embedding dimension *m* and scale to achieve optimal complexity assessment. Moreover, denoising across different scanners and individuals must find a balance between effectively removing noise and preserving the neural signals.

## Conclusion

In this study, we proposed a MVSE method incorporating T2* signals derived from 5T multi-echo EPI BOLD signals to improve the reliability of measuring neural activity complexity. The proposed method significantly improves the performance as compared to the conventional BOLD-based sample entropy approach. Notably, the method generated results in the ventricular regions that were more consistent with physical principles, and it showed superior performance in areas where ischemia with reduced neural activity. Moreover, when applied to both young and elderly cohorts, our method revealed significant differences across many brain regions. The proposed method enables a more precise and physically consistent analysis of neural activity complexity, thereby increasing the reliability of the entropy calculation.

## Supplementary Information

**SI Figure S1.**
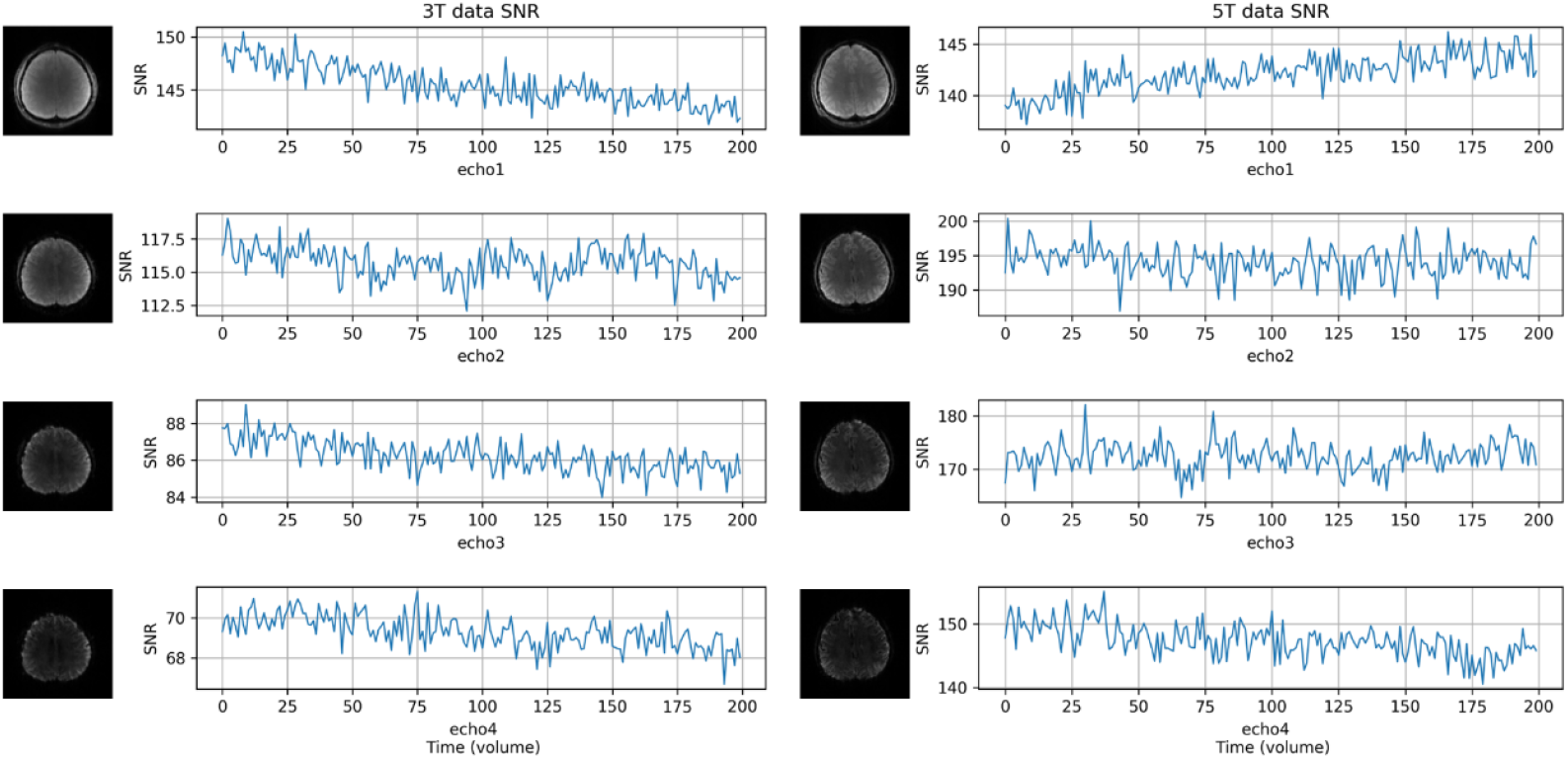
Temporal evolution of the signal-to-noise ratio (SNR) of the raw fMRI signals acquired at 3 T and 5 T.

**SI Figure S2.**
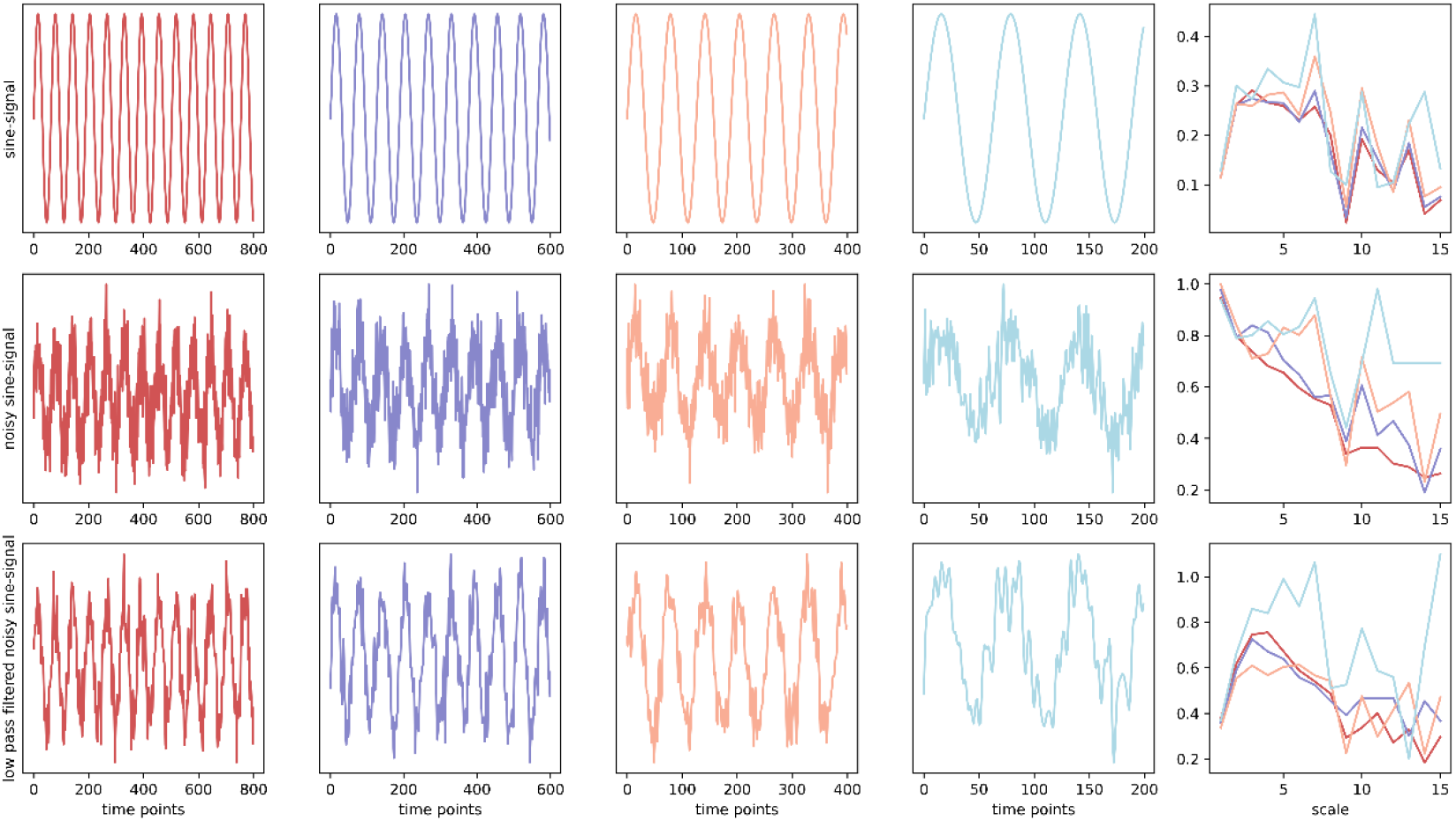
Variation of entropy across multiple scales for a single-frequency simulated signal with added noise

**SI Figure S3.**
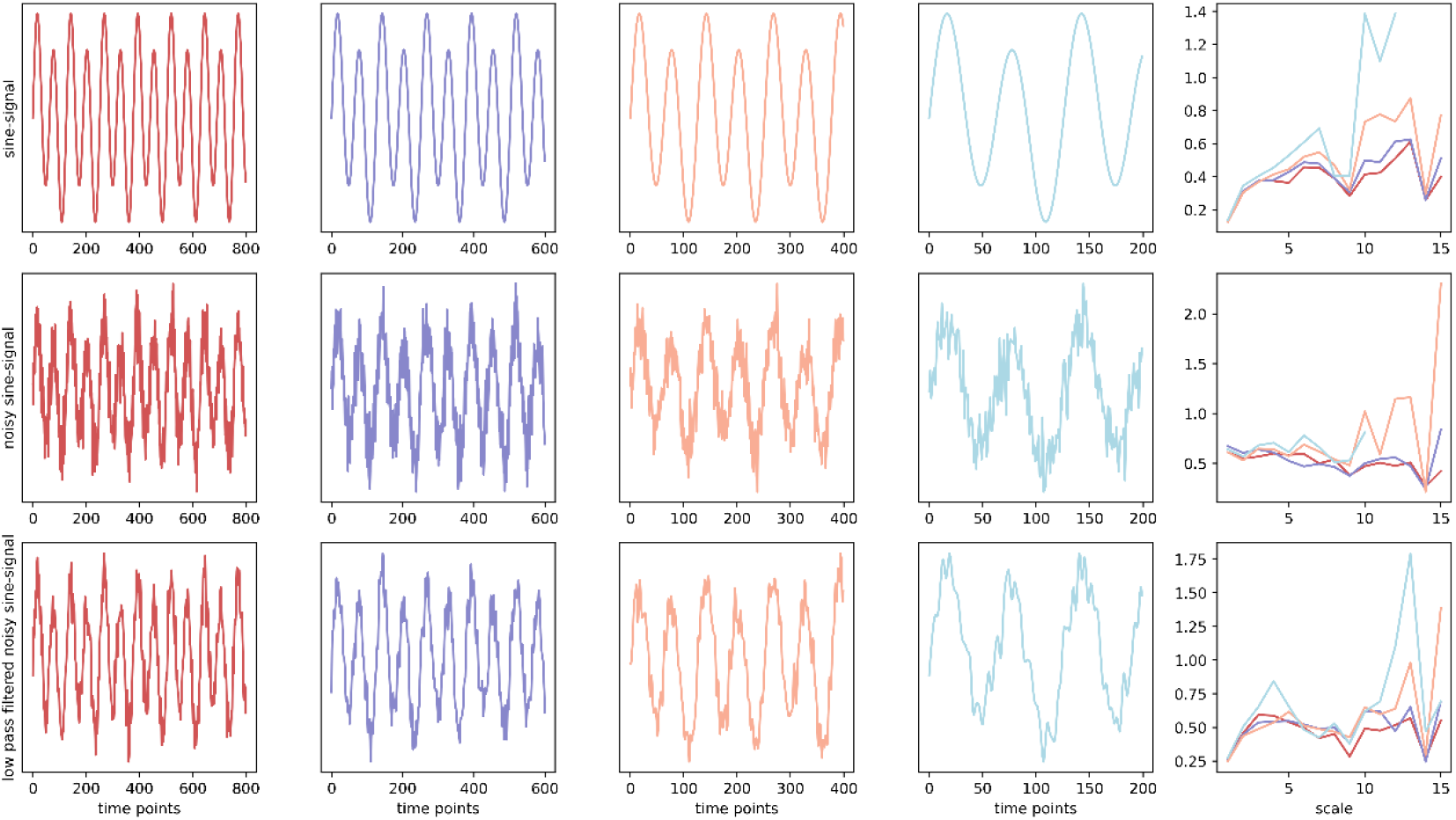
The variation of entropy across different scales for a multi-frequency simulated signal with added noise

**SI Figure S4.**
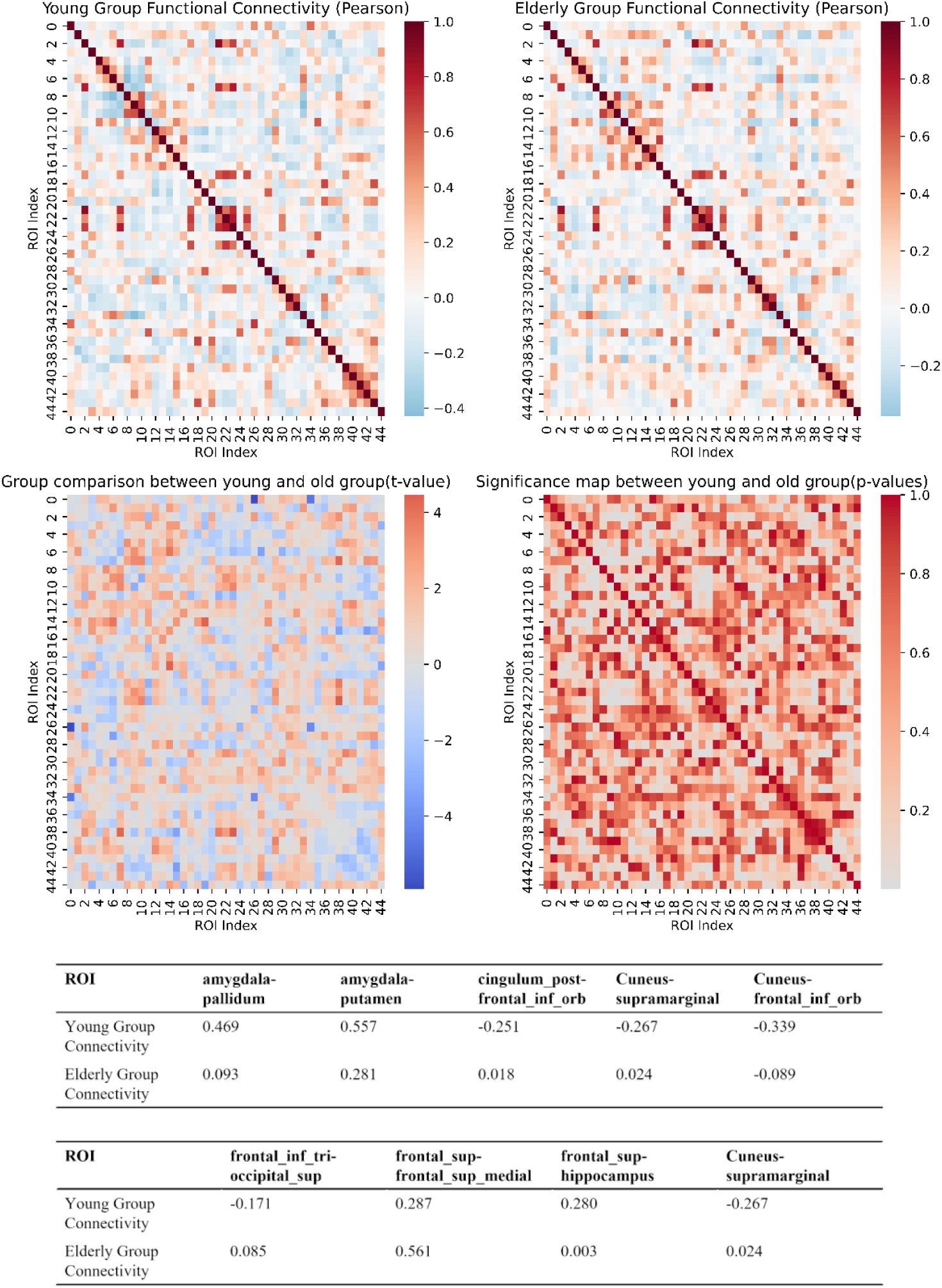
The functional connectivity across 45 regions of interest defined by the AAL atlas using Pearson correlation coefficients.

